# Shared neural codes for eye-gaze and valence

**DOI:** 10.1101/736462

**Authors:** Raviv Pryluk, Yosef Shohat, Anna Morozov, Dafna Friedman, Aryeh H. Taub, Rony Paz

## Abstract

The eye-gaze of others is a prominent social cue in primates and crucial for communication^1-7^, and atypical processing occurs in several conditions as autism-spectrum-disorder (ASD)^1,9-14^. The neural mechanisms that underlie eye-gaze remain vague, and it is still debated if these computations developed in dedicated neural circuits or shared with non-social elements. In many species, eye-gaze signals a threat and elicits anxiety, yet can also serve as a predictor for the outcome of the encounter: negative or positive^2,4,8^. Here, we hypothesized and find that neural codes overlap between eye-gaze and valence. Monkeys participated in a modified version of the human-intruder-test^8,15^ that includes direct and averted eye-gaze and interleaved with blocks of aversive and appetitive conditioning^16,17^. We find that single-neurons in the amygdala encode gaze^18^, whereas neurons in the anterior-cingulate-cortex encode the social context^19,20^ but not gaze. We identify a shared amygdala circuitry where neural responses to averted and direct gaze parallel the responses to appetitive and aversive value, correspondingly. Importantly, we distinguish two shared coding mechanisms: a shared-intensity scheme that is used for gaze and the unconditioned-stimulus, and a shared-activity scheme that is used for gaze and the conditioned-stimulus. The shared-intensity points to overlap in circuitry, whereas the shared-activity requires also correlated activity. Our results demonstrate that eye-gaze is coded as a signal of valence, yet also as the expected value of the interaction. The findings may suggest new insights into the mechanisms that underlie the malfunction of eye-gaze in ASD and the comorbidity with impaired social skills and anxiety.

## Main text

Recognizing and learning about potentially harmful or beneficial stimuli is crucial for survival of all organisms. In humans and primates in general, facial expressions, and in particular eye gaze of others, are a prominent and instructive signal ^2-7^. Averted or directed eye-gaze is a social signal that can indicate submissive vs. aggressive interactions, correspondingly. In agreement with this, eye-gaze was shown to elicit anxiety in primates ^4,8^, and evoke responses in the amygdala ^18,21-28^ – a brain region that serves as a hub for emotional responses in general and anxiety in particular ^25,29,30^. Moreover, gaze processing is disrupted in several neurodevelopmental and social disorders ^1,9-12^, and mainly in autism-spectrum-disorder (ASD) where abnormal activity of the amygdala is linked to gaze avoidance ^13,14^. However, eye-gaze is not only a valence-signal by itself, but can also serve as a predictor for future outcomes: aversive if an intruder makes direct eye-contact, or potentially rewarding if the intruder avoids eye-contact. This is in line with the amygdala not only playing a role in signaling outcome-valence (appetitive-aversive), but also in learning via affective conditioning ^16,17,31-34^ and signaling expectation for the outcome i.e. exhibit responses to a conditioned-stimulus (CS) ^16,33,35-38^. Yet it remains unknown whether similar mechanisms are used for coding of valence and eye-gaze; and moreover, whether a shared mechanism for coding of eye-gaze and outcome-expectation exists. Toward that end, we adapted the human intruder test (HIT) ^8,15,39^. HIT is widely used for assessing anxiety and defensive behaviors in non-human-primates, similar to the ‘stranger test’ in human infants ^40^. We performed recordings of single neurons during live interactions in a modified HIT paradigm that includes averted vs. direct eye-gaze of the intruder, and combined with an affective conditioning paradigm. We demonstrate that valence of both the outcome and its expectation are coded in similar networks as the eye-gaze of others, but via two different mechanisms.

Two monkeys participated in a modified version of the human intruder test (HIT) (Fig. 1a). Each HIT block consisted of 18 interactions with a human intruder that is seated behind an LCD shutter (<1ms RT) and either gazes directly at the monkey (eye contact, EC) or averted away from the monkey (no eye contact, NEC) when the shutter opens. These HIT blocks were interleaved with conditioning blocks of either appetitive or aversive trials (>=8 trials in a block, Fig.1b,Fig.1c), where the shutter opening serves as the conditioned stimulus (CS) and is followed by the outcome / unconditioned-stimulus (US, liquid reward or airpuff in appetitive/aversive blocks correspondingly) after a one second delay. We tracked the eye position of the monkeys and extracted four regions of interest (ROI): 1. the eye region of the intruder; 2. the face region of the intruder; 3. the whole shutter region; and 4. outside the shutter region (Fig.1d). Oculomotor behavior revealed distinct patterns (Fig.1d-g; Supp.Fig.1): shutter opening in the HIT blocks induced more interest in the eyes ROI compared to the conditioning blocks (Fig.1d, Kolmogorov-smirnov, p<1e-8, n-trials = 3108/1288 in HIT/conditioning trials; in 49 sessions, 24/25 per monkey). After exploring the eye of the human intruder, the monkeys continue to look more to the eyes/face ROI in blocks of direct gaze (Fig.1g, EC vs. NEC, *χ*2, p<1e-8, n-trial=1480/1628 in NEC/EC), similar to previous findings using static images of conspecifics^41^. We further aligned each trial according to the first time the monkey gazed at the intruder eyes (Interquartile range: 180-700ms) and found similar results (Supp.Fig.1, *χ*2, p<1e-6).

**Figure 1.**
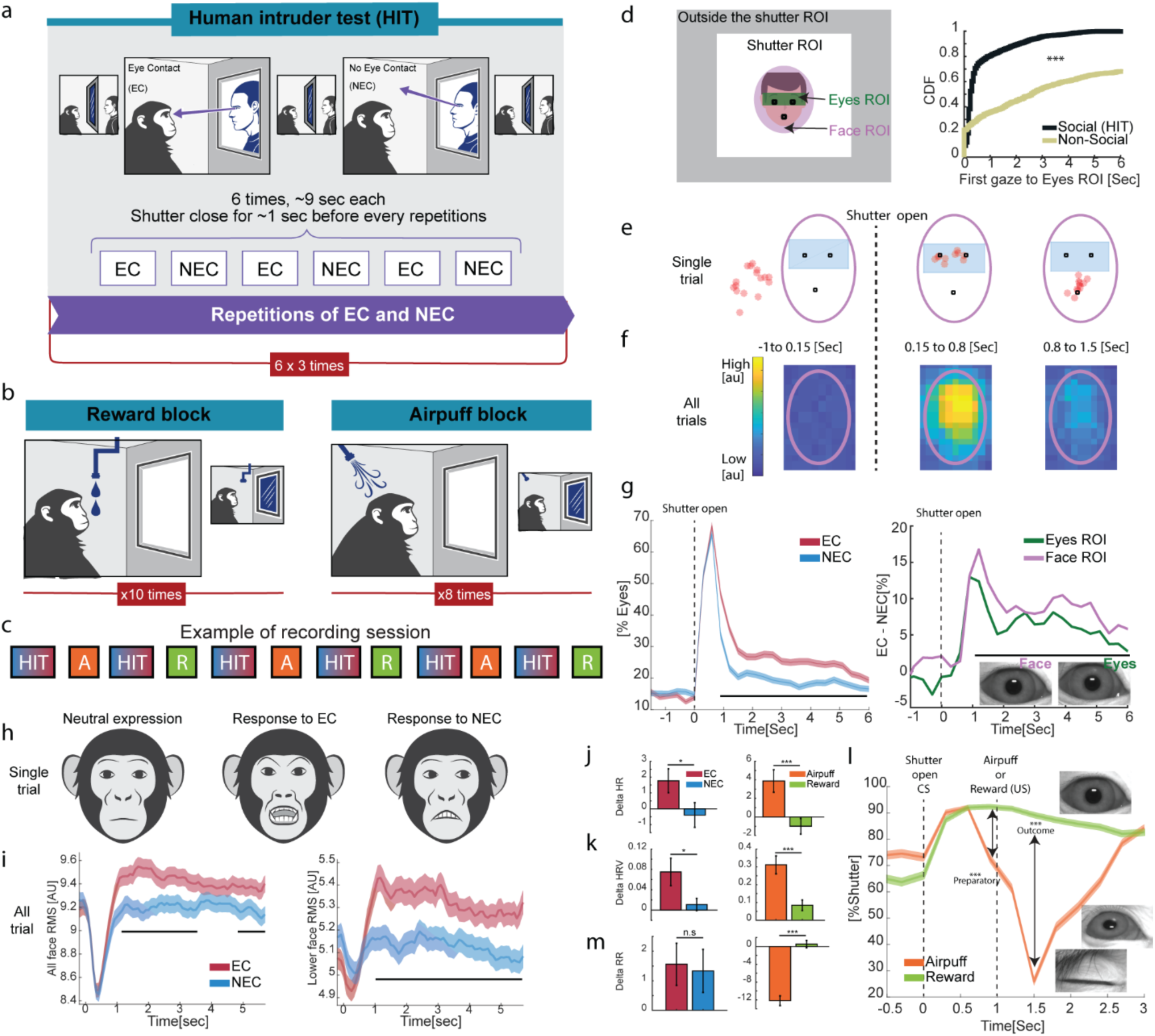
Paradigm and behavior during the Human-Intruder-Test (HIT) and affective conditioning blocks. a. Human Intruder Block: The shutter opens and closes 18 times, and the human intruder pseudorandomly alters between direct gaze (eye contact, EC) and averted eye-gaze (no eye contact, NEC). b. Classical conditioning blocks of either appetitive (reward) and aversive (airpuff). In these blocks, the shutter-open serves as a predictor (conditioned-stimulus, CS) to the appetitive/aversive outcome (unconditioned-stimulus, US). c. An example of the pseudorandom order of blocks in one recording session, with at least 120 seconds between blocks. d. Regions of interest (ROI) for the eye-tracking of the observer monkey (left): Only to the eyes of the human intruder (green); Only to the face of the intruder (pink); The whole shutter (white); The whole possible space (gray). Notice we report the same regions even when there is no intruder (the regions are similar across intruders because the faces are accurately aligned by positioning). e. Cumulative density function (right) of the first time after shutter opening that the monkeys look into the eyes ROI, separately for HIT and for conditioning blocks. f. Example of one shutter opening in an HIT block. Filled-circles (red) mark the location of the monkey’s gaze overlaid on an intruder real-position (schematic). Shown are three consecutive time windows (one before and two after). g. Density function of all eye locations in the HIT blocks in the same three consecutive time windows as in (E). Immediately after shutter opening and for few hundreds of milliseconds, the monkey looks mainly at the eyes of the human intruder. h. Left: Proportion of looking to the eyes-ROI in EC and NEC trials. The monkey first looks to the eyes region of the intruder, and then immediately breaks fixation in NEC trials significantly more than in EC trials. Right: The difference in proportion of the monkey’s look towards the eye and to the face ROI between EC and NEC trials. Both are significantly positive indicating that in EC trials the monkey maintains fixation to the face/eyes of the intruder. Upper black line represents a significant difference (p<0.05, *χ*2). i. Shown are schemes of typical facial expressions made by the monkeys in EC trials (middle, “aggressive”), in NEC trials (right,”interest”), compared to a neutral expression (left). See methods and Supp.Fig.2. j. The overall change in the facial expression in EC and NEC. Shown is the RMS of change in the image over the whole face (left) and only for the lower half of the face (right), compared to the neutral expression. Upper black line represents a significant difference (p<0.05, t-test). k. Differences in heart-rate between EC and NEC trials and between reward and airpuff trials. l. Differences in heart-rate-variability (HRV) between EC and NEC trials and between reward and airpuff trials. m. Response to aversive(airpuff) vs. appetitive(reward) in the oculomotor behavior n. Differences in respiratory-rate after shutter opens between EC and NEC trials and between reward and airpuff trials.

**Figure 2.**
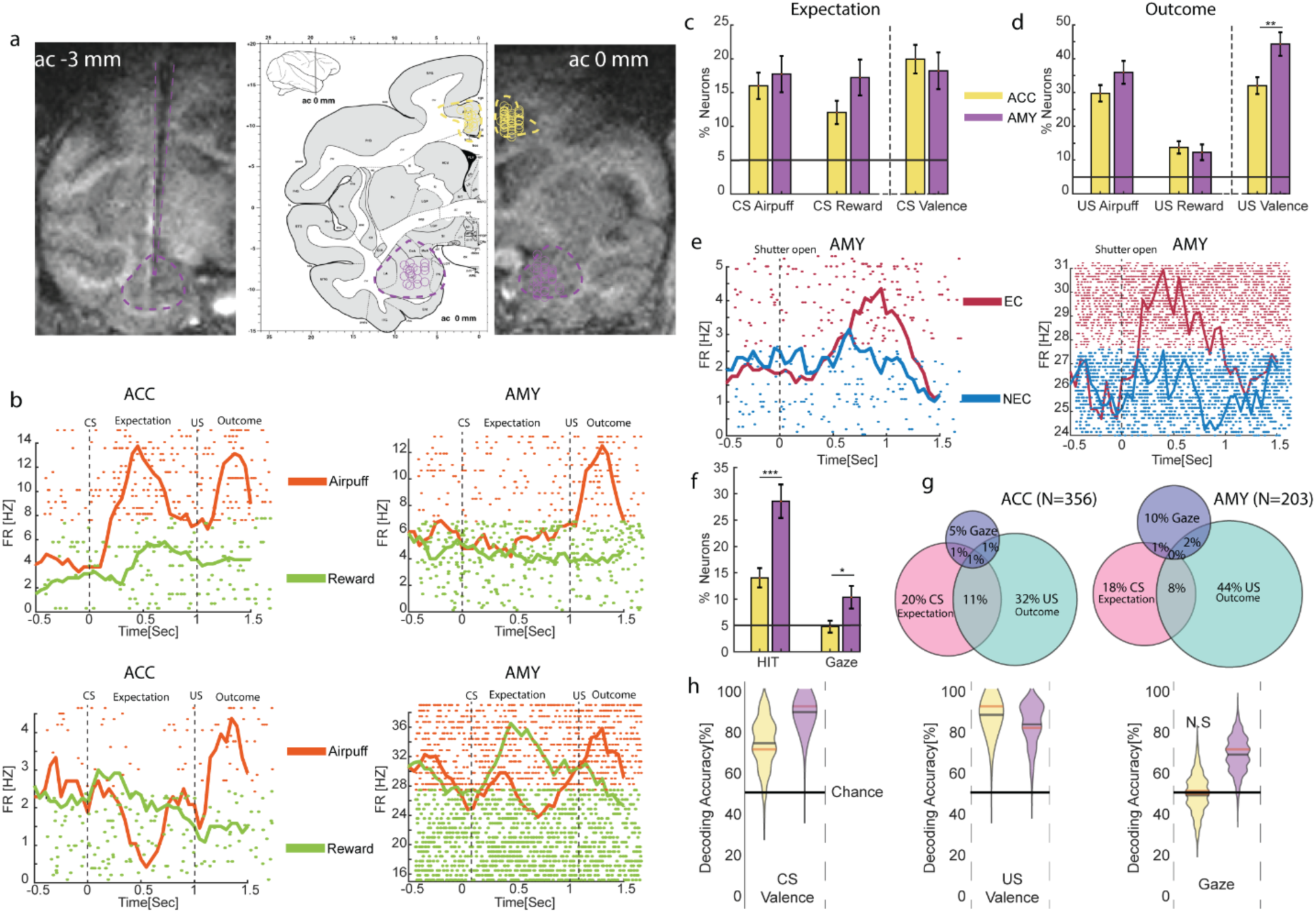
The amygdala codes for gaze and valence, and the ACC mainly codes valence. a. Recording locations: MRI with electrode directed into the BLA (AC=-3); Recording locations overlaid on a primate brain map (AC=0) and on an MRI scan (AC=0). b.PSTHs and raster plots of two representative neurons in the ACC and two in the amygdala during conditioning block. c. Proportion of neurons in the ACC and the Amygdala that respond to the CS (shutter open) in aversive trials (left), appetitive trials (middle), and discriminate between the two (right). d. Proportion of neurons in the ACC and in the amygdala that respond after the US (outcome) to the airpuff (left), reward (middle), and discriminate between the two (right). e. PSTH and raster plot of two representative neurons in the amygdala during the human-intruder block (HIT). f. The proportion of neurons in the ACC and the amygdala that respond significantly in the HIT blocks, in HIT blocks with eye-contact (EC), HIT blocks with no-eye-contact (NEC), and that discriminate between EC and NEC. g. Overlaps in the number of neurons that respond across the different tasks. The size of each area is proportional to the percentage of neurons. h. Population decoding accuracy for HIT vs. conditioning blocks. Discriminating appetitive from aversive with Amygdala and ACC neurons is significant using both the CS and the US responses, whereas only the Amygdala can decode gaze i.e. eye-contact from no-eye-contact.

We identified elicited facial expressions and found differences between EC to NEC trials. Specifically, the monkeys elicited more facial expressions when the intruders made eye-contact (Fig.1h,i, *χ*2, p<1e-2; Supp.Fig.2), in agreement with the stressful and defensive responses that are traditionally induced by direct-gaze of a human intruder ^8^. Heart-rate and heart-rate-variability (HRV) further confirmed anxiety-related responses ^42^ (Fig.1j,k, t-test, p<0.05).

In the conditioning blocks, the monkeys quickly learned to distinguish and anticipate the different outcomes (appetitive / aversive) after shutter opening in each specific block (Fig.1l, *χ*2, p<1e-3). In aversive-airpuff blocks, they closed the eyes after shutter opening both in preparation for the airpuff as well as immediately after its delivery (Fig.1l, *χ*2, p<1e-3). In addition, they held their inhale before the expected airpuff but not before reward (Fig.1m, t-test, p<1e-3).

In conclusion, there was a clear differential behavioral response in HIT sessions between eye-contact of the intruder and no-eye-contact, and there was a clear differential response between appetitive and aversive blocks, for both the unconditioned-stimulus (US, outcome), and the conditioned-stimulus (CS, preparatory/expectation).

To examine and compare neural responses, we recorded single units from the basolateral-complex of the amygdala (BLA) and the anterior-cingulate-cortex (ACC) (Fig.2a, n=24/25 sessions per monkey, n = 356/203 neurons in the ACC/Amygdala, 224/103 and 132/100 per monkey). In the conditioning blocks, we define two epochs: a preparatory/expectation epoch (CS-related, after the shutter opening but before US delivery), and an outcome epoch (US-related, following delivery of airpuff / reward) (Fig.2b). Confirming previous studies, we find that neurons in the amygdala and the ACC respond to the appetitive CS (Amy: 35/203, ACC: 43/356, *χ*2, p<1e-3 for both), to the aversive CS (Amy: 36/203, ACC: 57/356, *χ*2, p<1e-3 for both), and also discriminate between valence (Amy: 37/203, ACC: 71/356, *χ*2, p<1e-3 for both). Moreover, similar proportions of cells were responsive in the two regions (Fig.2c, 2, p>0.09 for all). Similarly, neurons in the ACC and in the amygdala responded to the appetitive US (Amy: 25/203, ACC: 49/356, p<1e-2 for both), and aversive US (Amy: 73/203, ACC: 106/356, p<1e-3 for both). Here again there were similar proportions in both regions (*χ*2, p>0.1). However, more amygdala neurons discriminated valence i.e. between appetitive and aversive outcome (Fig.2d, Amy: 90/203, ACC:114/356, *χ*2, p<1e-2).

In the HIT blocks (Fig.2e), neural responses were computed from the epoch when the monkey first looks to the eyes-ROI, namely this is the first time the monkey can differentiate if it is an EC or an NEC trial (Interquartile range: 180-700ms, see Fig.1d). There were much more responsive neurons in the amygdala than in the ACC during HIT blocks (Fig.2f, Amy: 58/203, ACC: 50/356, 2, p<1e-3), and more amygdala neurons discriminate between EC and NEC of the intruder (Fig.2f, Amy: 21/203, ACC: 17/356, *χ*2, p<0.05). The number of ACC neurons that coded for the intruder gaze was not different than chance (Binomial test, p>0.1). We tested for overlap in responses and found that the proportion of neurons that responded to both gaze and valence was not different than chance, both in the amygdala and in the ACC (Fig.2g Binomial test, p>0.1).

We conclude that at the single cell level, valence is coded in the ACC, whereas both gaze and valence are coded in amygdala neurons.

We noticed that the proportion of amygdala neurons that code for gaze is low compared to the proportion of neurons that code for valence, both for CS-related responses and for US-related responses (Fig.2c,d,f, *χ*2, CS: p<0.05, US: p<1e-3). This finding is in line with previous studies in both monkeys and humans ^18,43^. However, because a neuron can contribute at the population level even if it does not exhibit a significant response by itself ^44^, we further tested whether the population of recorded neurons holds information about the eye gaze of others by training a linear decoder on population vectors ^45,46^. In accordance with the single-cell analyses, population activity in the amygdala and the ACC could discriminate between appetitive and aversive trials, both using CS-related and using US-related activity (Fig.2h, bootstrap analysis with CI 95%). However, only the amygdala population could discriminate between EC and NEC trials, whereas the ACC population did not exceed chance-level (Fig.2h, bootstrap analysis with CI 95%).

To conclude, both at the single-cell and at the population level, the two regions code for valence, whereas only the amygdala codes for the eye-gaze of the intruder.

The finding that the amygdala holds information about valence and eye gaze of others within the same circuitry suggests that there might be a shared coding mechanism via shared neural ensembles. In order to test this hypothesis of shared coding for valence and gaze, we used the decoder approach again, but this time we trained on one type of trials and tested on another. If discrimination accuracy is above chance-level, this would mean that the population uses similar mechanisms to hold information for one situation - appetitive vs. aversive, as for the other - EC vs. NEC. We therefore trained a linear decoder to distinguish between trials of EC and NEC and tested it on distinguishing between trials of aversive and appetitive. Importantly, this was done separately for the CS-related and the US-related responses.

In agreement with the aforementioned finding that the ACC does not hold information about eye-gaze, the decoding performance was not different than chance in both CS and US related activity (Fig.3a,b top insets, bootstrap analysis with CI 95%). In contrast, performance was significantly above chance level when using amygdala population, and moreover, it was the case when using both CS-related activity and US-related activity (Fig.3a,b, Supp.Fig.3, bootstrap analysis with CI 95%). Performance was approximately linear in the number of neurons, starting from chance-level and rising to more than 80% accuracy when using all available amygdala neurons (CS: 82.5%, US: 80%, n=203, p<0.001 for both) (Fig.3a,b bottom insets). This linearity demonstrates that the shared coding of valence and eye gaze is not due to the few neurons that had significant responses to both contexts (Fig.2g), a notion that was further supported by the finding that accuracy remained highly similar when dropping these few neurons (CS: 81%, n=201; US: 79%, n=198).

These findings demonstrate that a shared coding mechanism is used by amygdala neurons, because the decoder was trained only on gaze discrimination, yet successfully tested on valence discrimination.

In general, there could be two shared activity patterns that would allow training on one context and decoding the other. In the first, termed here *shared-activity*, the same neurons respond similarly to gaze and valence (Fig.3c). This means that the same neurons respond in the same direction and with similar proportion (decrease/increase firing rates proportionally) for NEC vs. EC as for appetitive vs. aversive. Namely, a neuron’s response is correlated along eye-gaze and valence. Alternatively, in the second option termed *shared-intensity*, shared populations of neurons respond only in the same direction (high or low firing-rates) to gaze and valence, yet individual neurons are not correlated across the contexts (Fig. 3d). We therefore tested which mechanism applies here, and is it different when using CS-related or US-related epochs. To do so, we applied several different tests.

**Figure 3.**
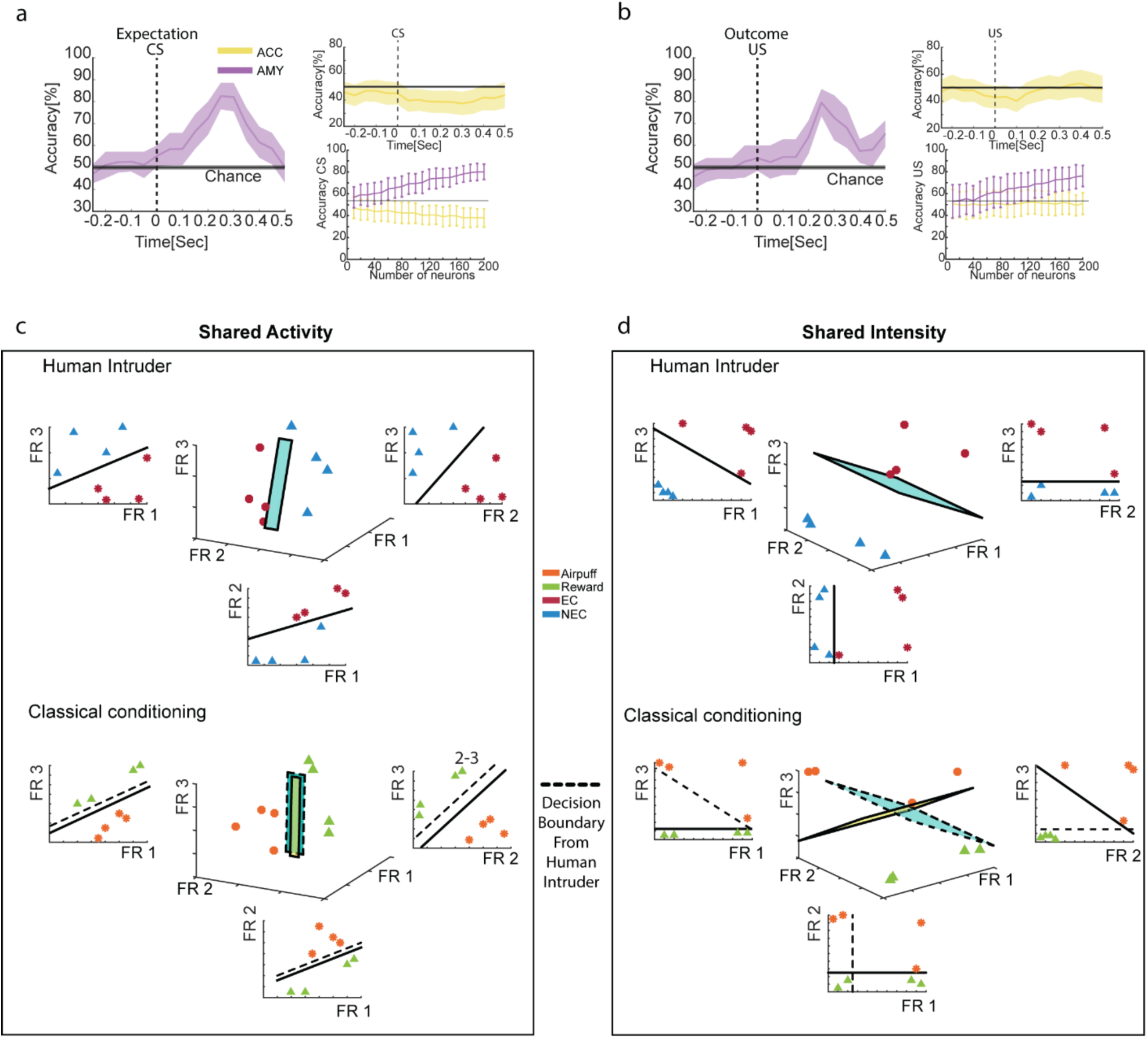
Shared coding for valence and gaze in amygdala neurons. a. Population decoding accuracy when training on eye-gaze (EC vs. NEC) and testing on valence (aversive vs. appetitive), using CS-related activity. Right-top inset: Similar format using ACC population. Right-bottom inset: Peak decoding accuracy using increasing numbers of neurons (Mean +/-STD, bootsrap). b. Same as (A) but using US-related activity. c. A scheme in 3D demonstrating the *shared-activity* mechanism. Here, neurons respond similarly to gaze and valence, meaning their response is correlated across NEC-to-EC and Appetitive-to-Aversive. The optimal linear separator computed for the HIT between each pair of neurons is shown in solid line, and the surface computed for 3 neurons is shown in the middle. A similar presentation is shown for the conditioning, but with the linear separators from the HIT imposed in dashed lines. For *Shared-activity*, they are more parallel than perpendicular. Therefore, the decoding is highly similar when training on one context and testing on the other d. A scheme in 3D demonstrating the *shared-intensity* mechanism. Here, different neurons in the population respond with similar changes in firing rate to gaze and valence, but individual neurons are not correlated across gaze and valence. For *Shared-Intensity*, the linear separators imposed from the HIT on the conditioning are less parallel and more perpendicular. However, neurons provide enough spikes for an overall firing-rate to allow correct decoding.

**Figure 4.**
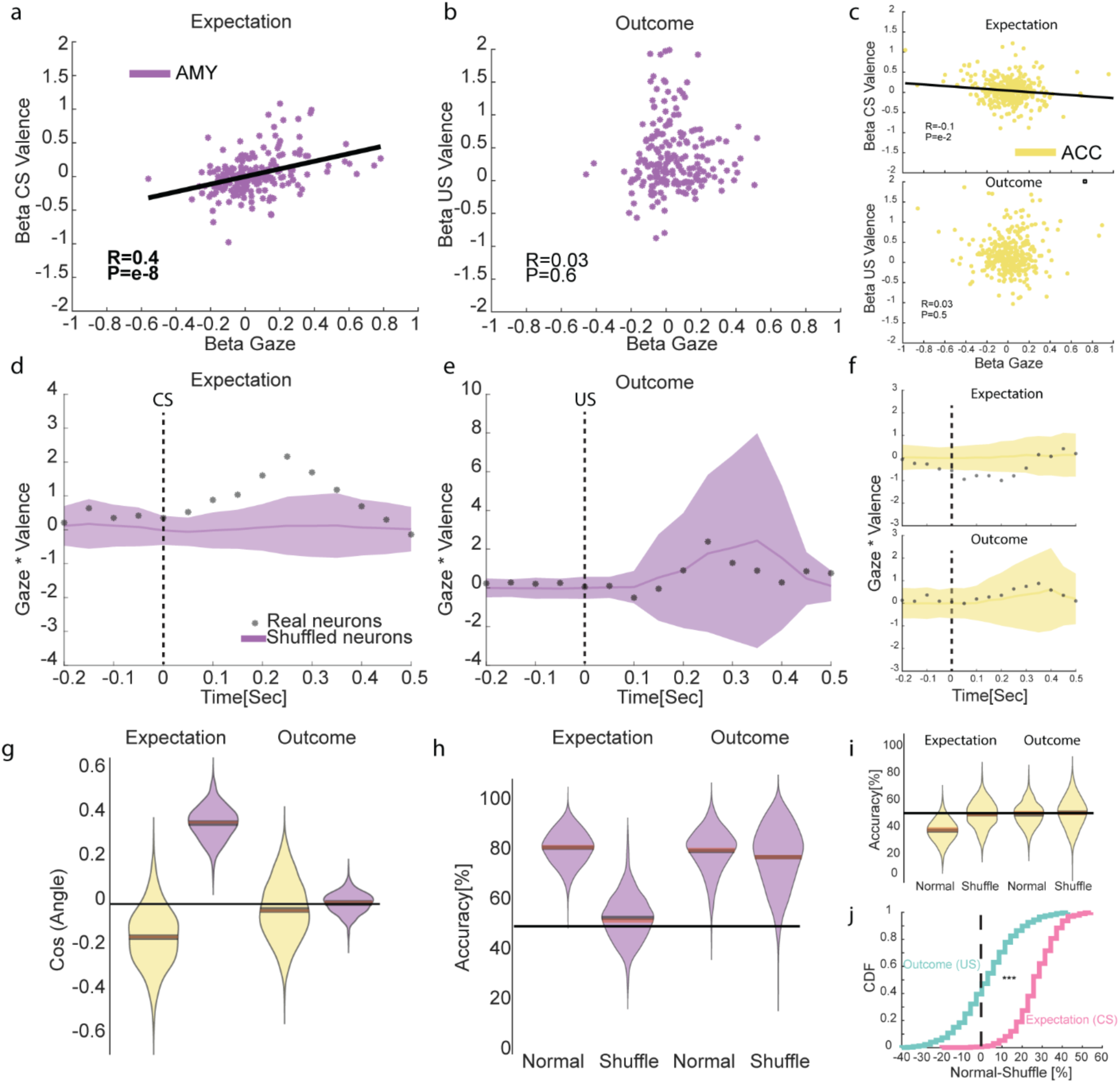
A *shared-intensity* coding for eye-gaze and US-valence and a *shared-activity* coding for eye-gaze and CS-valence. a. Correlation between the linear-regression coefficients of gaze (eye-contact vs. no-eye-contact, x-axis) and of valence (aversive vs. appetitive, y-axis) using CS-related activity. All amygdala neurons are shown. b. Same as (A) using US-related activity. c. Same as (A) and (B) for ACC activity, CS-related (top) and US-related (bottom). d. Neurons respond in the same direction for eye-gaze and valence using CS-related activity (scalar-product between the coefficients of gaze and of valence for each neuron). Black asterisks represent data from real neurons and shaded-magenta is 95% confidence interval based on bootstrap shuffle. e. Same as (D) using US-related activity. f. Same as (C) and (D) for ACC activity, CS-related (top) and US-related (bottom). g. The angle between the decision boundaries derived from the population-vector of gaze and valence separately (shown is the scalar product between the two vectors). h. Population decoding accuracy for real and shuffled neurons using CS-related activity. i. Same as (H) for ACC activity. j. Cumulative-distribution of the difference in decoding accuracy between real and shuffled neurons.

We first examined the two shared coding options at the single cell level, and used linear-regression on the neural responses for gaze (comparing EC to NEC) and separately on the responses for valence (comparing aversive to appetitive). We obtained and compared two separate coefficients: *β*_*valence*_ that represents the difference in firing rate between airpuff and reward, and *β*_*gaze*_ that represents the difference in firing rate between EC and NEC. If the two coefficients are similar for individual neurons, it means the neurons code valence and eye-gaze not only along the same direction but also with similar modulation proportion (reward-airpuff as NEC-EC). We found that the two coefficients were positively correlated in amygdala neurons, but only when using CS-related activity and not when using US-related activity (Fig.4a-c, Pearson correlation, amygdala: CS: r=0.4, p<1e-8, US: r=0.03, p>0.5; ACC: CS: r=-0.1, p<1e-2, US: r=0.03, p>0.5). This observation supports a *shared-activity* between valence and gaze for the CS, yet a *shared-intensity* for the US epoch. The *shared-intensity* in the US is further supported by direct examination of overall increases/decreases in firing-rates for direct gaze and US valence (Supp.Fig.4, Z-test p<1e-3).

This finding was further validated by examining the scalar product between the two coefficients (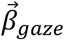 and 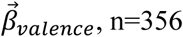, n=356 in the ACC and n=203 in the amygdala). If more neurons respond in similar proportion (*shared-activity)*, then the scalar-product would be positive; otherwise, the scalar product will be close to zero if neurons respond in random order (or negative if in opposite directions). In the amygdala, using CS-related activity outperforms a shuffling test (Fig.4d, bootstrap across neurons), yet using US-related activity it does not (Fig.4e, bootstrap test). In the ACC, neurons were similar or lower than the shuffled test (Fig.4f, bootstrap test). In addition, the mean value for the US-related shuffled activity is higher than for the CS-related shuffled activity (Fig.4d,e, CS=0.1, US=1.8, bootstrap, p<0.05). This is because more neurons both in gaze and in US-valence increase their firing rate, resulting in a higher positive scalar product for shuffled neurons, further supporting the *shared-intensity*. In contrast, in the CS the similarity in the response increases the scalar product in the actual neurons but not in the shuffled population.

To test the two shared coding mechanisms at the population level, we computed the angles between the decision boundaries of the two linear decoders: one boundary that separates EC from NEC and one that separates aversive from appetitive. When computed over the US epoch, or using ACC population, the decision boundaries of valence and gaze are not different than perpendicular (dot-product not significantly different from zero), whereas only using CS activity from the amygdala population shows a significant difference from perpendicular decision boundaries (Fig.4g, bootstrap analysis CI 95%). Finally, we trained the decoder on gaze and tested on valence while shuffling the order of neurons. This approach is used to test if it is the specific ensemble of neurons that matters, or just an overall increase in firing rate. In line with the previous results, performance using amygdala activity from the CS epoch was decreased dramatically when comparing actual to shuffled neurons (Fig.4h,Supp.Fig.5 bootstrap analysis with CI 95%), whereas using US activity it even slightly increased (Fig.4h,i,j, Supp.Fig.5 bootstrap analysis with CI 95%), further supporting the two different shared mechanisms: *shared-activity* between gaze and CS-valence, and *shared-intensity* between gaze and US-valence.

The eyes of others became a more prominent signal along evolution due to anatomical changes in facial morphology that have forced a shift in salience from the shape of the face to the eyes ^2^. The importance of the amygdala in processing of eye-gaze was shown in humans and in macaques ^13,18,23,47-51^. Here, we recorded neural activity in the amygdala and the ACC during live interactions in a modified version of the human intruder test (HIT) that includes also a conditioning paradigm. Whereas both regions differentiated between valence in their CS-related and US-related responses ^32^, only the amygdala differentiated between averted vs. directed gaze of an intruder. Importantly, in the amygdala, both the CS-related and the US-related responses were shared with the eye-gaze and in a valence-specific manner, namely appetitive (reward) to aversive (airpuff) paralleled averted-to direct-gaze. Our results obtained in live-interactions, across aversive-to-appetitive valence axis, and for eye-gaze, suggest that social value evolved from or in parallel to primary-reinforcer value. Therefore, processing of social stimuli and among it gaze does not necessarily occur in dedicated neural circuits ^2,52-54^.

Importantly, we identified two different shared mechanisms that allow decoding of value based on responses to eye-gaze. Both shared mechanisms had the same directionality on the valence axis, namely appetitive to aversive match averted to direct gaze, both used the same neural circuitry, but were different in implementation. The *shared-intensity* which is shared across gaze and outcome (US) occurs by an overall increase in firing rate. This is a more ‘primitive’ mechanism that points to shared origins in the same circuit, where an aversive outcome is similar in value to a predator gaze^4^ or to a threat by a peer. This is also in line with the findings of the human-intruder-test where gaze elicits immediate anxiety^8^. The coding of expectation, i.e. a learned CS, was also shared with eye-gaze responses; however, it was shared via a *shared-activity* mechanism that in addition required that the responses would be correlated on a single-neuron basis (rather than only on average over the population). Because this is a more demanding mechanism and because the amygdala has evolved in parallel to the development of social interactions ^55-57^, we suggest that these responses later enabled complex social processes such as learning by observation also mediated by the amygdala ^12,58,59^ and social-based decision-making in extended circuits ^5,19,60,61^. Specifically, it can be used to anticipate social outcomes based on context, a direct gaze likely calls for a challenge and hence predicts danger and a coming attack; whereas an averted gaze usually predicts a submissive and permissive encounter and potentially rewarding (mating, food sharing/offering).

The shared coding we identify here between value, eye-gaze, and social-dependent expectation can explain the complexity of several neurodevelopmental disorders and especially ASD, where simple aspects of gaze processing are disrupted^1,9-14^ and higher social skills as understanding of other intentions, emotions, and actions are all compromised and accompanied with abnormal amygdala function^2,62-64^. Finally, it suggests a direct mechanism for the co-morbidity between ASD and anxiety-disorders, and might open a window to novel therapeutic interventions in amygdala circuits.

## Methods and Supplementary Figures

### Methods

#### Animals and surgical procedures

Two male macaca fascicularis (4–8 kg) were implanted with a round recording chamber above the amygdala and ACC covering both regions in both hemispheres. All procedures were approved and conducted in accordance with the regulations of the Weizmann Institute Animal Care and Use Committee (IACUC), following NIH regulations and with AAALAC accreditation.

MRI based electrode positioning scans were acquired twice, on a 3-Tesla MRI scanner (MAGNETOM Trio, Siemens) with a CP knee coil (Siemens) and using 0.53mm resolution. A first scan before surgery used to align and refine anatomical maps for individual animals (relative location of the amygdala, ACC and anatomical markers such as the interaural line and the anterior commissure), and to guide the positioning of the chamber on the skull. After surgery, we performed scans with deep electrodes directed toward the amygdala and the ACC (see for example Fig.2a), and calculate the anatomical anterior– posterior and lateral-medial borders relative to the penetrations. The depth of the amygdala is calculated from the dura surface.

#### Electrophysiology recordings

Each day, 1-3 multichannel (16 contacts each) microelectrodes vector arrays (NeuroNexus) were lowered into the brain using an electrode-positioning-system (NAN, Israel). Vectors were moved independently into the amygdala and ACC while identifying electro-physiological markers tracking the known anatomical pathway. We allowed 30 min for the tissue and signal to stabilize before starting acquisition and behavioral protocol. Data is pre-amplified and stored at 22Khz for later processing. In real-time a 0.3Hz-6KHz band-pass filter and on-line spike sorting is performed using a template-based algorithm (Alpha Lab Pro, Alpha Omega). Off-line spike sorting was performed on the raw data for all sessions to improve unit isolation (offline sorter, Plexon Inc).

#### Behavioral Paradigms

Fast LCD shutter (307 × 407 mm) is placed between the monkey and the intruder (FOS-307 × 406-PSCT-LV; Liquid Crystal Technologies) to block visual site. Direct current (48v) through the LCD shutter turns it clear/transparent with an onset/offset rise time of <1ms. To enhance precision for neural activity we place a photodiode (BPX65 Silicon PIN Photodiode) that can be detected with onset/offset of <1e-4ms. There are three types of blocks in each daily session: Human intruder; Airpuffs; Liquid rewards. The blocks are randomized along a session, with more than 120 seconds separating blocks (Fig.1).

##### Human Intruder

Each block includes 6*3 shutter openings, in which the human intruder alters between Eye-Contact (EC) and No-Eye-contact (NEC) in a pseudorandom order. In both EC and NEC the human maintains gaze direction for 6-9 secs independently of the monkeys’ behavior. We generated a per-day pre-defined sequence of EC and NEC with 3 options of sequences that alter across sessions: seq1 (BlockA: EC,NEC,EC,NEC,EC,NEC; BlockB: EC,EC,NEC,NEC,EC,EC; BlockC: NEC,NEC,EC,EC,NEC,EC); seq2 (BlockB, BlockC, BlockA) and seq3 (BlockC, BlockA, BlockB). This was aimed to randomize and prevent learning of EC/NEC order, but also to provide across-days statistics for neural recordings. The human intruder face was filmed and all the trials were monitored to validate that the intruders indeed maintained constant gaze and followed the daily sequence.

##### Reward

Each block contains 10 trials with an inter-trial-interval of a pseudorandom 20-40 secs. In each trial the shutter opening serves as the conditioned-stimulus (CS) and was followed after 1sec delay by few drops of juice delivered to the monkey’s mouth.

Airpuff - Each block contains 8 trials with an inter-trial-interval of a pseudorandom 20-40 secs. In each trial the shutter opening serves as the conditioned-stimulus (CS) and was followed after 1sec delay by air puff (5-15 Psi; located 5 cm from the face).

The monkeys had information about which block is about to start as the human intruder paradigm starts with 5 secs of pure sinus wave (300Hz) followed by the human intruder entering the room and sitting in front of the monkey, with closed shutter. The monkey could not see any part of the human unless the shutter is open.

#### Behavioral analysis

##### Eye tracking

A stationary monocular eye tracker was installed for the purpose of eye tracking and gaze estimation. The system included two cameras (Ximea_MQ013RG) – one for eye capturing of the monkey and one for intruders’ monitoring and an infrared LED light bar (MetaBright Exolight ISO-14-IRN-24) for face illumination and corneal reflection (CR) production. The eye-recording camera efficiently captured the CR due to its near IR (infra-red) property.

Software implementation was based on the open source project ‘OpenEyes’ ^1^, which allows the estimation of subject’s point of gaze (POG) on the field of view (FOV) projection. In our case, the FOV scene images were extracted from the video stream of the intruder monitoring camera. The ‘OpenEyes’ framework makes use of the Starburst algorithm ^2^ for finding the pupil contour, and assesses the POG by the means of pupil center and CR method ^3^. The conditions of our experimental setup (brightly lighted room, large CR of near-rectangular shape and brown sclera of the subject ^4^) required a slight modification of the original algorithm for pupil and CR detection. In our variation of the software, the shot noise reduction was skipped, and the CR wasn’t removed from the image after its detection, due to its large size. To find the pupil center, we extended the Starburst algorithm. After finding the features candidates for pupil contour, instead of fitting ellipse using RANSAC (random sample consensus) paradigm, we used the “imfindcircles” Matlab function, which searches for circle-candidates applying Hough transform based algorithm. To generate the input for the function, edges image was produced by gradient magnitude calculation followed by binarization. This procedure resulted in a black image with white edges, and was passed to “imfindcircles” with object polarity parameter set to “dark” (specifying that the object – the pupil - is darker than its background). The function returns a list of candidate circles, ordered by circle strengths. Starting from the circle with the biggest strength, the list is searched for the first circle containing a predefined number of minimum feature points that were extracted by the Srtarburst algorithm. Finally, the pupil center is estimated by the center of the found circle. A standard calibration procedure was performed, whereby the monkeys sequentially fixated on 3×3 known grid points in the scene image (according to the original openEyes implementation). To cause the subject’s fixation, the screen with the shutter closed, was consecutively illuminated by a laser pointer in the 9 locations. The exact frames of subject’s fixation were detected and coordinated with the illumination timings (each time the laser is activated, it records the exact time in the system). The human intruders were filmed throughout all the interactions with the monkeys, and their faces and eyes were marked both automatically and manually for validation. The 9 (3×3) fixation points were filmed by the same camera, allowing the projections of the fixation points and the intruders on the same plane. Each frame from the eyes of the monkeys therefore result in a point (x and y position) on this plane, allowing to calculate the gaze of the monkey in one of the four ROI’s – eyes of the intruders, face of the intruders, shutter region and all the rest.

##### Facial expression

One Ximea_MQ013RG camera filmed the face region of the recoded monkey in 34Hz. For every recording session, the mean image during the ‘alone’ period was calculated (i.e. when the monkey was alone in the room with closed shutter). This mean image (See Supp.Fig.2) was subtracted from every frame taken during the Human Intruder interactions. Root Mean Square (RMS) of all the pixels in this subtracted frame is then calculated and the mean and STD are presented for EC and NEC trials (Fig.1i). Additionally, each day we manually define 3 ROI’s – upper face, lower face and ears (See Supp.Fig.2). The same analysis is repeated separately to each ROI and differences between EC and NEC were validated across both for upper and lower face (Fig.1i and Supp.Fig.2)

##### Heart Rate and Respiratory rate measures

Piezoelectric pulse transducer: The cardiac and respiratory traces (for measure of Heart-rate, Heart-rate-variability and Respiratory-rate) ^5,6^ were recorded using a piezoelectric pulse transducer (UFI, model 1010) in 2790Hz. We use an elastic belt about 23cm (9 inches) long and fasten extender belt to one end of transducer package using VELCRO™ closures all wrapped around the monkey’s chest. We use a piezoelectric pulse transducer (UFI, model 1010) glued around the center allowing direct sensing the heart pulse.

For validation, the respiratory trace is recorded also using solid-state transducer which measures changes in chest or abdominal circumference due to respiration (UFI, model 1132) at 2790Hz. The signal from the piezo sensor also provides respiratory rate parameters, allowing two independent measures for comparison and calibration of parameters.

The piezo-electric signal was processed using a custom made Matlab software. A respiratory signal\trace was extracted using a first order Butterworth filter, and smoothed with running windows. Respiratory peaks were then extracted using ‘findpeaks’ function. A cardiac signal\trace was extracted by subtracting the filtered respiratory signal from the raw piezo-electric signal. The resulting signal was then processed for each day separately, using filtering and findpeaks parameters. The parameters of the day-specific processing were derived by comparing different sets of parameters to manually tagged cardiac peaks from each day. The resulting day-tailored processed signal was validated using manual inspection of all trials. In addition, the quality of each trial was manually rated, and noisy signal epochs were marked to validate that the result is not due to trials of insufficient quality.

Respiratory rate and heart rate measurements were calculated for each trial using a sliding window of 1 second and heart rate variability (HRV) using running window of 5 seconds, yielding a continuous signal for further analysis. The HRV measure is the standard deviation of normal-normal beat interval (SDNN), a well-established and frequently used measure ^7^. Finally, we normalized the changes in each measure by subtracting the mean value from the closed shutter epoch before each trial, to obtain evoked responses.

#### Neural activity analysis

##### Single neuron analysis

The analysis of the neural data focused on three times epochs. In the human intruder blocks, we focused on 400-700ms after shutter opening. This time was chosen because of the oculomotor behavior of the monkeys (Fig.1) showing that the first time it can identify if it is an EC or an NEC trial has an Interquartile range of 180-700ms (see Fig.1d for the full CDF). All analyses were repeated (see Supp.Fig.1, Supp.Fig.5) also when aligning each trial according to the actual time in that trial that the EC/NEC information is available (first gaze to eyes ROI). Such an alignment was done in order to focus on the differences between EC and NEC of the intruder and because fixation shape neural activity ^8,9^. In the affective (reward/aversive) conditioning blocks, the neural data was taken from 0-300ms after the conditioned-stimulus, termed CS-related activity; and from 0-300ms after reward/airpuff delivery (outcome), termed US-related activity.

Neural activity is normalized according to the baseline activity before the relevant block, using the same window length (300ms) to calculate the mean and standard deviation of the firing rate. Therefore, the normalized (z-scored) firing rate is:

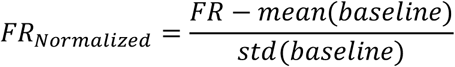

These z-scores were used to quantify the percentage of responsive neurons to the different stimuli. T-tests are used to compare valence (airpuff to reward) or gaze (EC to NEC), and chi-square or binomial tests are used to compare proportions of neurons.

##### Population decoding

Pseudo-simultaneous population response vector is used for the decoding analysis. The same procedure as reported in details ^10,11^ is used. The population vector contains spike counts of each neuron in a specific time bin. Each brain area has its own vectors, and the number of vectors is defined by the number of available trials:

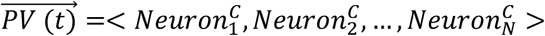

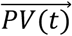 is the response vector of specific trial in condition C, in time bin (t), in a brain region that has N neurons. We use the same number of neurons in the amygdala and ACC, therefore we randomly discarded excess neurons in the ACC, resulting in 203 neurons in both.

There are four conditions, airpuff and reward that belong to the valence class and EC and NEC that belong to the gaze class. In the analysis that was conducted in Fig.2 we trained and tested within the same class, whereas in all other analyses we trained on one class and tested on the other class. If we change the order in the training, such that training for NEC yield airpuff and training for EC yield reward, the decoding accuracy is exactly (100-CorectDecoding, see Supp.Fig.3). For both the training and testing we used linear classifier based on maximization procedure of the SVM algorithm (fitSVM Matlab function). Each training set yield a boundary line (set of weights for every neuron) and a threshold that separates the two conditions under consideration. The same output from the training was then used to assess the accuracy in the test set.

For a given neuron and a given condition we used 80% of the trials for training and 20% for testing when done within the same class. When we trained on one class and tested on the other, we used all the available trials for training and testing. The accuracy of every decoder was estimated by pseudorandom resampling from the available trials 1,000 times.

In the analysis of Fig.4 we shuffled the neurons such that the index of each neuron in 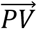 is randomly assigned. Therefore, the spike count of every neuron remains, but its position in the vector changes.

##### Decision boundary analysis

In order to estimate if the mechanism that allows decoding of one class based on the other is due to shared-activity or shared-intensity, we estimate the angle between the boundary lines. Every training sample yields a vector of weights:

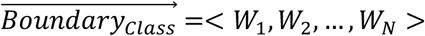

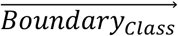 is the decision boundary of one training sample in a brain region with N neurons. Every brain region has two boundaries, one for gaze and one for valence.

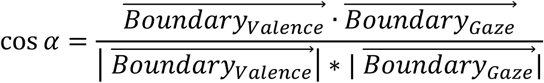

Each of the boundaries is sampled 1,000 times to obtain for every sample of the two boundaries an angle. The results are presented as cos(*α*) and not *α*, so zero (0) values represent perpendicular boundaries.

##### Linear regression analysis

We estimated the tuning of the neurons to valence and gaze by linear regression analysis. The firing rate, FR, of every neuron is fitted during every time bin with one of the following equations:

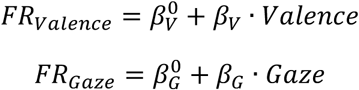

*Valence* is 1 for airpuff trials and −1 for reward trials, whereas *Gaze* is 1 for EC and −1 for NEC. The regression analysis yield for every neuron two coefficients, *β*_*V*_ and *β*_*G*_.

##### Scalar product of linear regression coefficients

We calculated the scalar product between 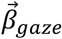 and 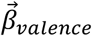 where the vector sign indicates it is a vector of all neurons in a certain brain region 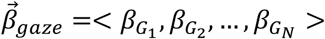 and 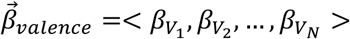The intuition behind this scalar product is that if more neurons response in a similar direction, than it is expected to be positive and vice versa.

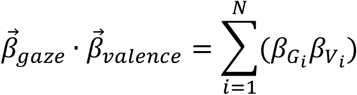

We also calculated a shuffled version where a random index is used, and hence the multiplication of the coefficients is done across two different neurons. The shuffled scalar product is repeated 1,000 times.

## Acknowledgments

We thank Dr. Yoav Kfir for scientific and technical advice; Drs. Eilat Kahana and Nir Samuel for medical and surgical procedures; Daniel Goldin for engineering design, Dr. Edna Furman-Haran and Fanny Attar for MRI procedures. This work was supported by ISF #2352/19 and ERC-2016-CoG #724910 grants to R. Paz.

---

**Supp.Fig.1.**
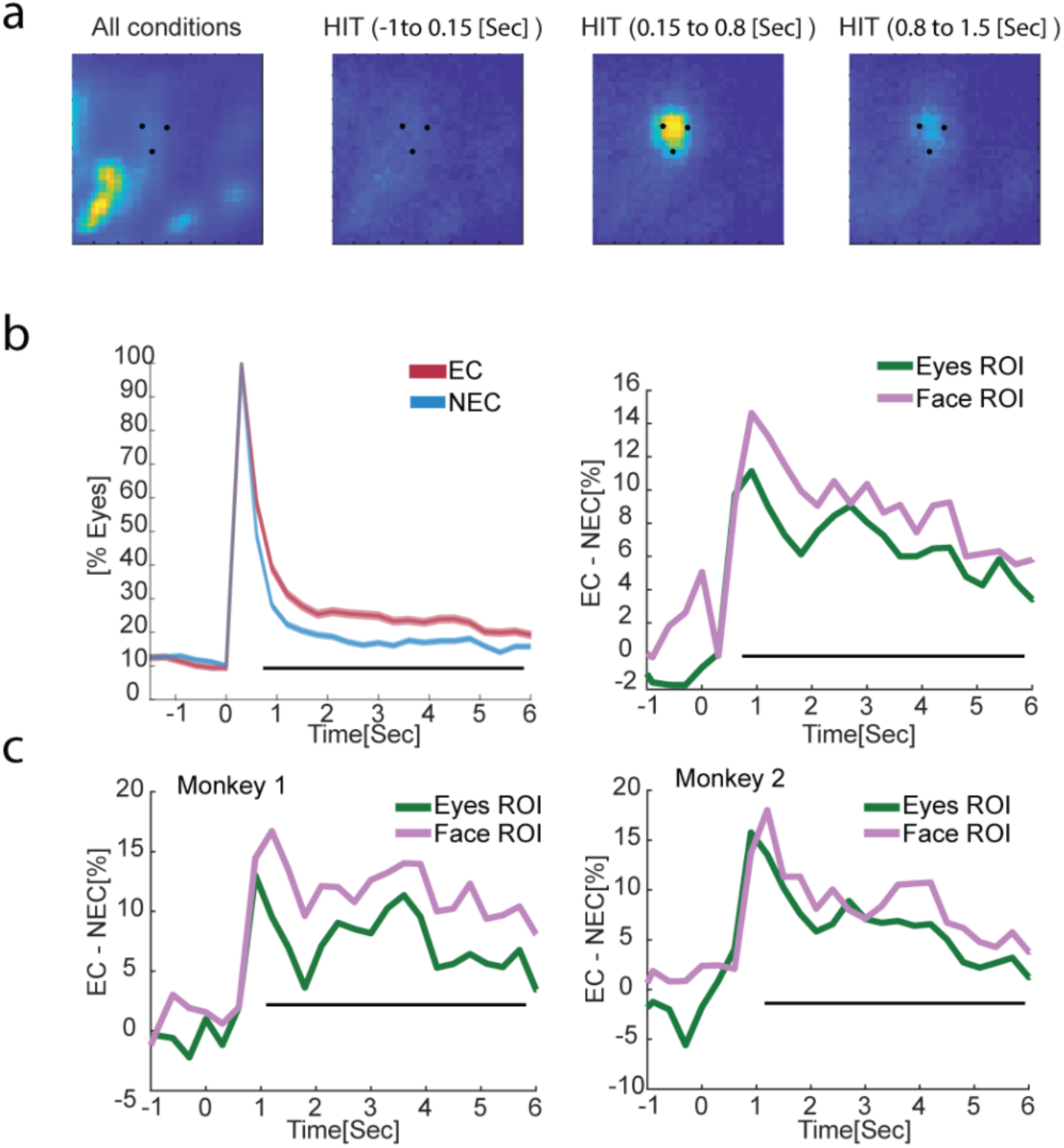
Differential behavioral response to EC and NEC. a. Same format as Fig.1f but for all shutter ROI (and not just face ROI). As can be seen, the monkeys look at the face and eyes ROI mainly in the human intruder interactions. b. Same format as Fig.1g, but aligned to the first time the monkeys looked to the intruder’s eyes ROI in each trial separately. c. Same format as Fig.1g-right, separately for each monkey

**Supp.Fig.2.**
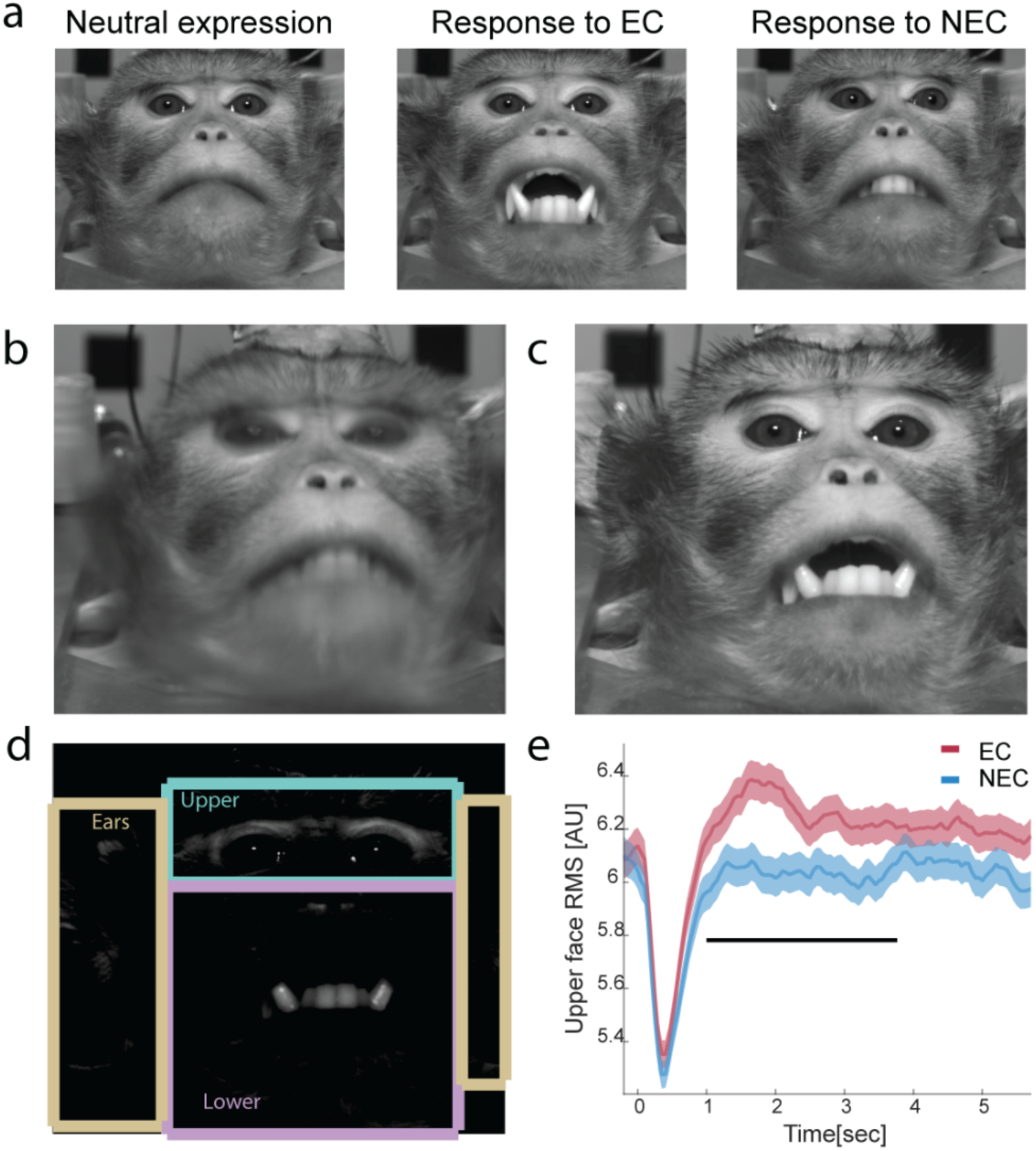
Extracting differences in facial expression. a. Same as Fig.1h depicting the actual faces. b. Shown is the mean image computed from all frames taken from before the Human Intruder Interaction (i.e. when the monkey is alone with closed shutter). c. An example of a frame during EC (eye contact) interaction. d. The mean frame (b) is subtracted from every frame during the interaction (i.e. as in b), to obtain a ‘diff’ image. Three ROIs are defined manually for every day – Upper, Ears and Lower. e. Root Mean Square of every ROI is calculated. Shown are differences between EC and NEC in the Upper part (see main Fig.1 for other parts).

**Supp.Fig.3.**
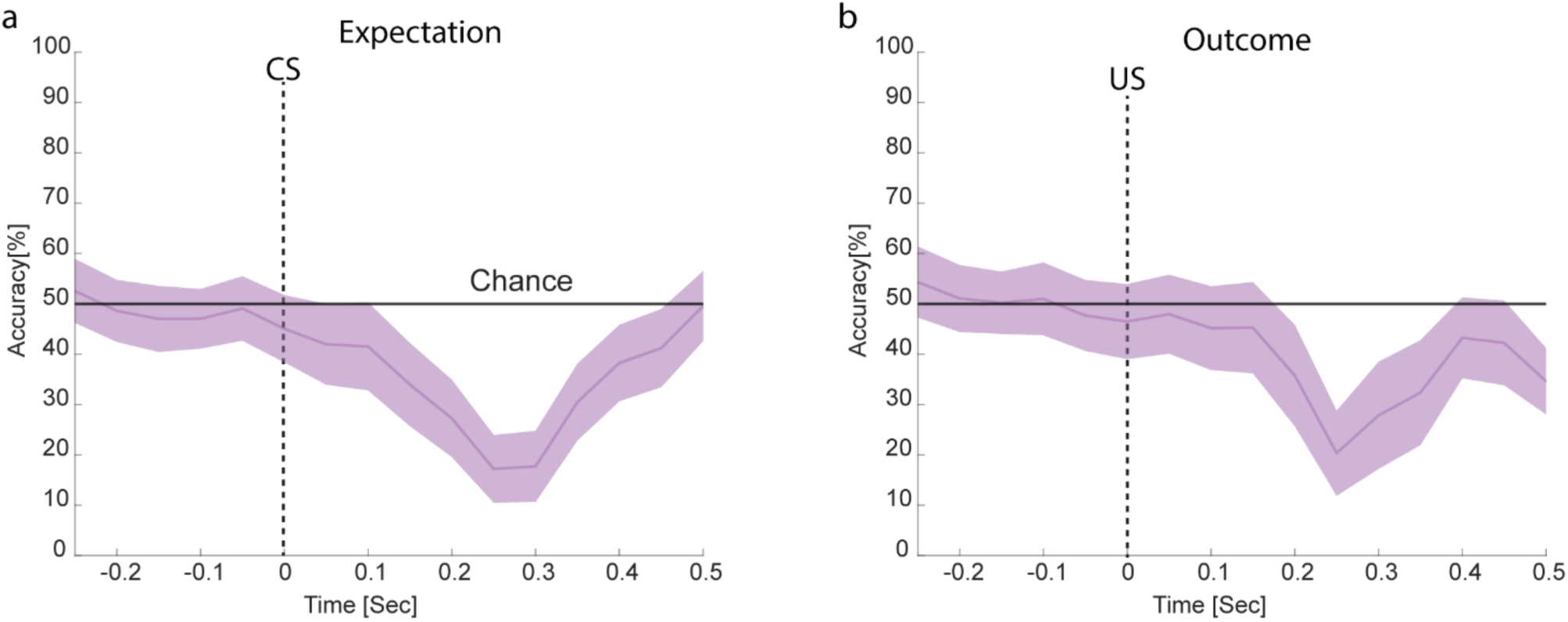
Decoding when reversing valence. a. Same format as in main Fig.3a,b. Population decoding accuracy but when training on eye-gaze (NEC vs. EC) and testing on valence (aversive vs. appetitive), using CS-related activity. b. Same as (a) but using US-related activity.

**Supp.Fig.4.**
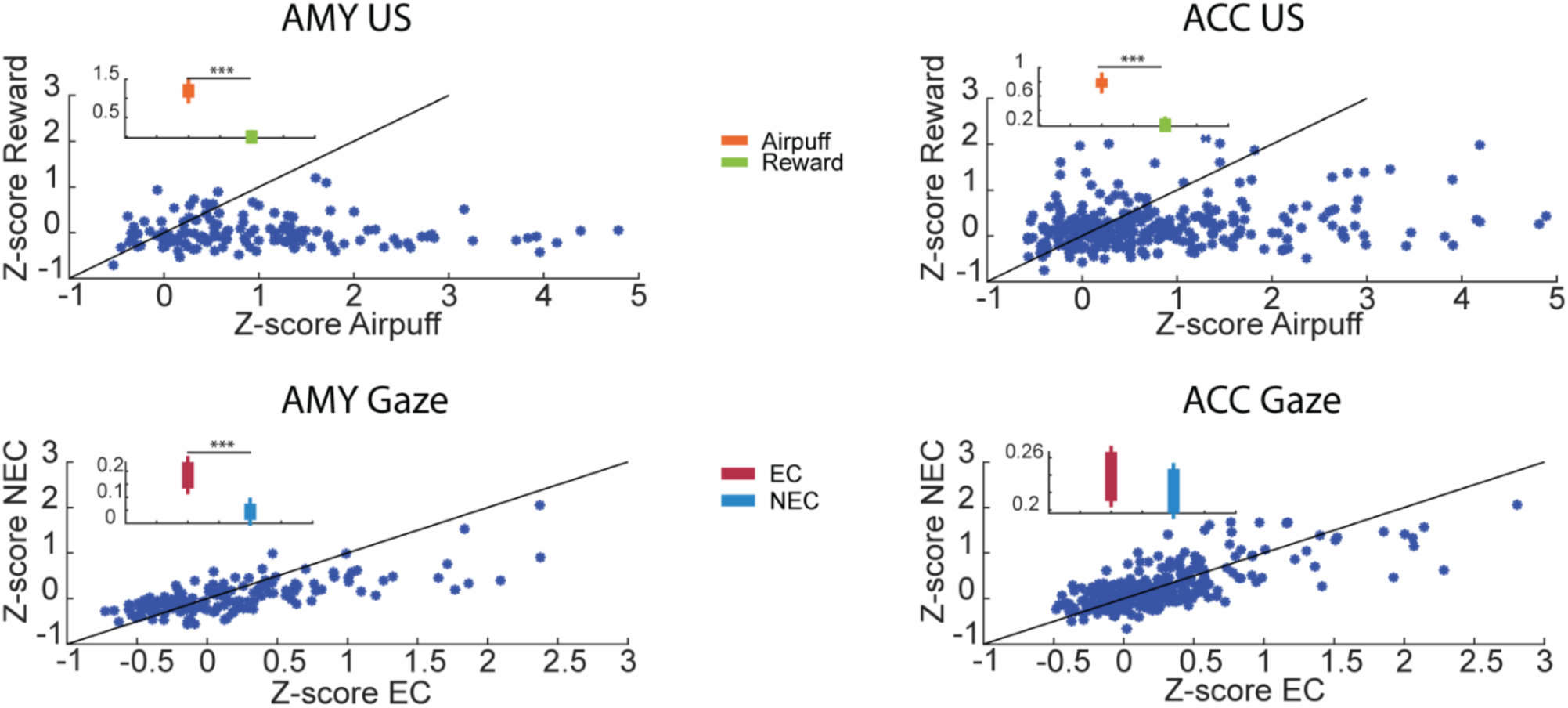
Single-neurons activity compared across conditions. If *shared-intensity* drives the successful decoding in the US epoch, we expect to find an overall change in the firing rate (increase or decrease) for gaze and for US valence. Indeed, we find that there are more valence positive neurons (increased firing rate to airpuff) in the amygdala in the US epoch, and that there are more gaze positive neurons (increased firing rate to EC) in the amygdala.

**Supp.Fig.5.**
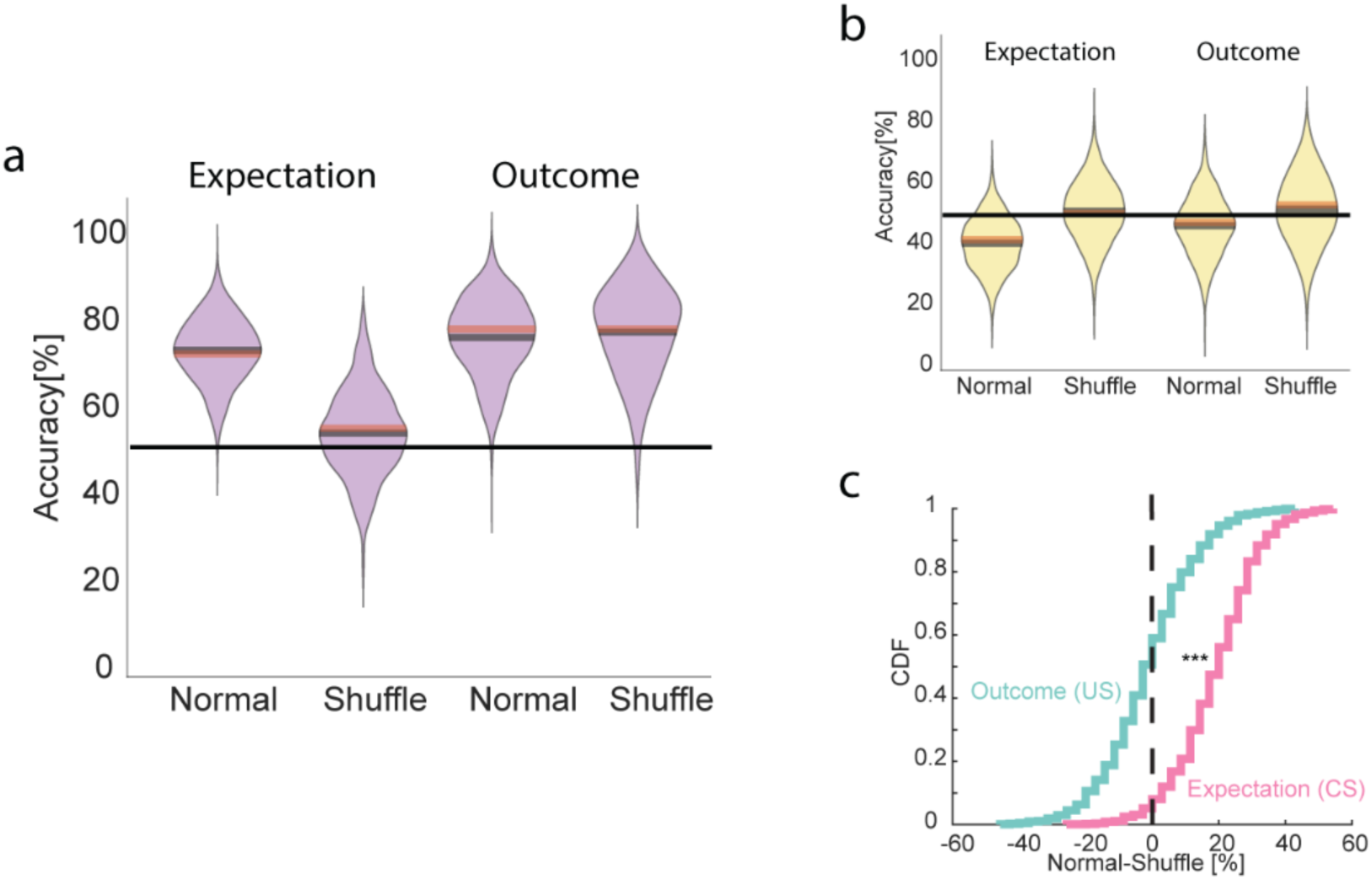
Decoding with alignment to Shutter opening (each trial separately) a. Same format as in Fig.4h,i,j. Population decoding accuracy for real and shuffled Amygdala neurons. b. Same as (a) for ACC activity. Cumulative-distribution of the difference in decoding accuracy between real and shuffled neurons.

